# Validation and optimisation of wearable accelerometer data pre-processing for digital measure implementation and development

**DOI:** 10.64898/2026.03.21.713324

**Authors:** Joss Langford, Jia Ying Chua, Ian Long, Alan C. Williams, Melvyn Hillsdon

## Abstract

The increasing use of accelerometers as digital health technologies in clinical trials and clinical care is driving the need for data processing to meet medical standards. The aim of this study was to create and test a modular pipeline for the pre-processing of high-resolution accelerometry that assures the quality, transparency and traceability of digital measures from sensor-level data. The objective is for the pipeline to be a foundational layer in the development, implementation and comparison of measures.

The study developed the open GENEAcore package to meet the requirements of regulators, verifying the engineering implementation and analytically validating outputs against reference datasets. Early stages included the optimisation of calibration and non-wear detection. Data-driven detection of behavioural transitions was then validated to give direct bout outputs without the need to identify rules for epoch aggregation and interruptions. The utility for measure development was shown by comparing two algorithms for the characterisation of activity intensity in both the epoch and bout paradigms.

Non-wear was detected with a balanced accuracy of 92.3% and the commonly used 13mg acceleration standard deviation threshold was empirically validated for the first time. The detection of transitions proved reliable with 99% detected, on average, within 2 seconds of their occurrence to give a mean expected event duration of 68.6s from a log-normal distribution. The different activity intensity algorithms were more than 99% concordant during movement but their outputs diverged in low movement conditions. Importantly, variable duration bouts created 31% higher daily activity durations compared to 1-second epochs.

This evaluation of pre-processing steps has confirmed the attention to detail required to create robust and reproducible results for later clinical validation where small changes in an algorithm or its implementation may have clinically meaningful consequences.

## Introduction

### Overview

Public health and physical behaviour research have been significantly advanced by the advent of wearable raw data accelerometers [1]. Increasingly, these sensor-level measurement systems are used in commercial research to evaluate medical products as digital health technologies [2] and to guide clinical care [3]. In contrast to consumer-grade devices with embedded algorithms, the original measurement outputs are preserved, allowing full verification of the sensor engineering [4] as well as futureproofing for new algorithms.

Wearable sensors make it possible to collect high-resolution information about an individual’s physiology, behaviour and daily routines over long periods with low burden to participants and healthcare professionals. Unlike traditional episodic measures (for example, from clinic visits), these data are gathered continuously in real life. They can be more sensitive, timely, and representative of someone’s complete health and behaviour [5]. As well as reducing measurement error (misclassification), these objectively recorded data eliminate the recall and desirability response biases that come with relying on people’s memory, subjective judgement or self-report [6]. These biases, particularly recall [7] and desirability [8], can lead to over- or under-estimates of time in active behaviours that in turn attenuates associations with health outcomes [9].

However, challenges remain in processing the large volumes of data from objective measurement into digital health measures that are meaningful to both patients and clinicians [10]. To be trusted, digital measures need to be accurate, reliable and clearly associated with health outcomes in the clinical evidence [11]. They also need to be produced through a process that is transparent and repeatable with an understanding of any potential risk to safety [12]. Regulators also insist on full traceability of the intended use throughout the development, implementation and operation of the system [13].

As raw data technologies move from research settings into clinical applications, there is growing pressure to make the way we handle data both more efficient and more trustworthy. Clear and consistent pre-processing, where raw sensor data are prepared before further analysis, is then an essential foundation [14]. Standardised pre-processing strengthens interoperability between devices and facilitates reproducibility of algorithm implementation. It lowers costs, speeds up development, and lays the groundwork for regulatory approval, which in turn supports clinical adoption and reimbursement.

In this work, we describe and test a pre-processing framework for wearable accelerometer measurements (Fig 1).

**Fig 1.**
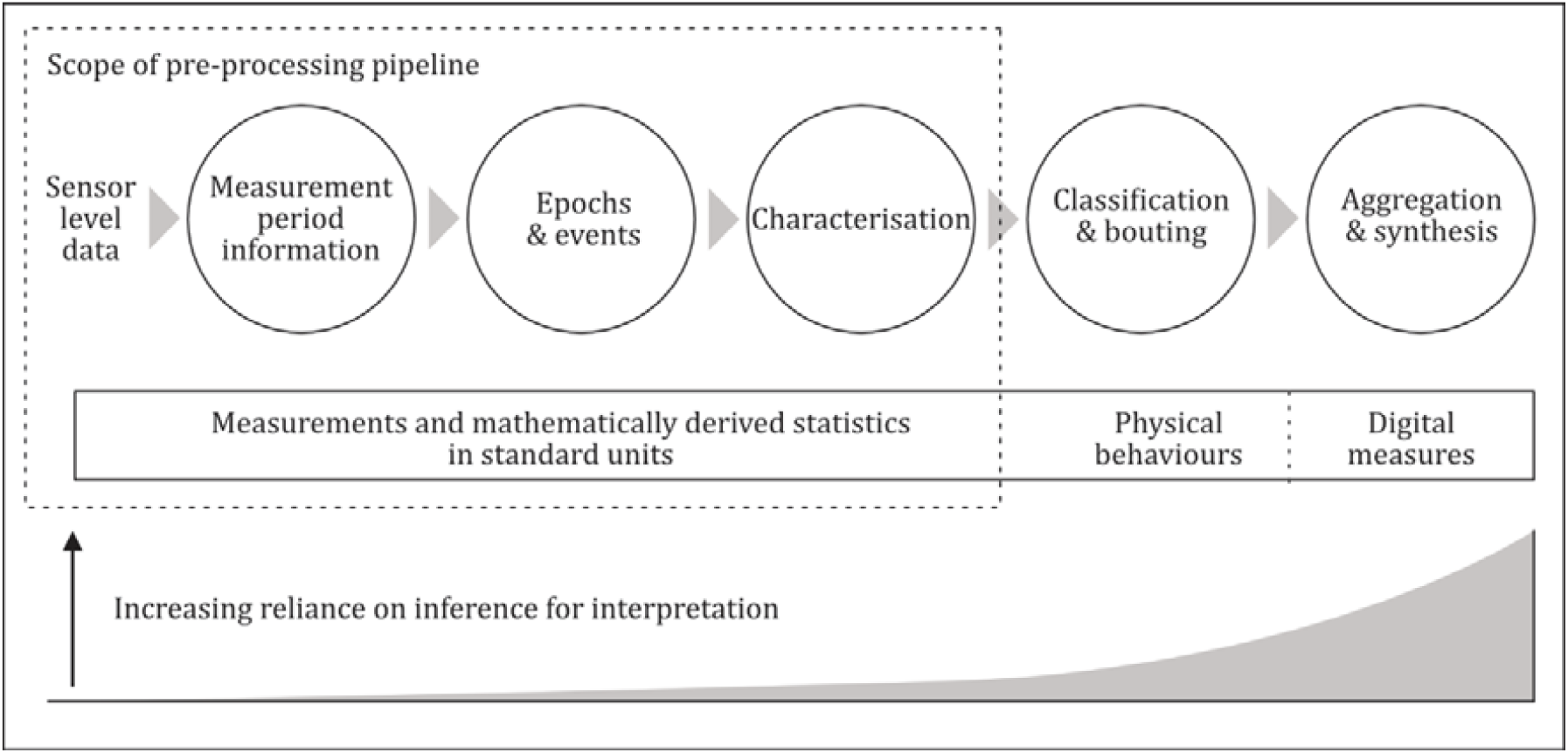
Generic sensor data processing pipeline. A representation of a generic sensor data processing pipeline showing the scope and aims of this work in the context of classified physical behaviours, digital measures and the level of inference required for interpretation.

The framework uses open-source software in an integrated test environment to extend verification from the original device measurements through to the digital measures output from the system. It is a fast, flexible layer for the implementation, assessment and analytical validation of digital measures.

The scope of this work covers the first 3 stages of Fig 1, transforming raw sensor data into contiguous periods of pre-processed data, ready for classification into physical behaviours (for example, sleep or walking). Our objective in this early processing is to keep assumptions to a minimum, and to document them carefully when they are required. We want to create a transparent, reproducible and data-driven process to give the strongest possible foundation for more complex later stages. While both heuristic and machine learning approaches are essential tools for analysing digital measures in the context of health outcomes, the acceleration measurements directly represent real-world behaviours and standard units (SI) can often be preserved for reproducibility and interoperability.

The overall aim of the study is to define an open, modular pipeline (GENEAcore) for the pre-processing of high-resolution accelerometry that can assure the quality and traceability of raw, sensor-level data as they are successively refined into digital measures. The process must be transparent and efficiently deliver essential summary information concerning data source integrity, interoperability and quality. This manuscript will document the verification of the engineering implementation and the analytical validation of outputs against reference datasets.

### Pre-processing pipeline

There are several examples of processing pipelines for accelerometry data openly available for use by researchers. Some rely on heuristic assumptions (GGIR [15]) or large machine learning datasets (OxWearables [16]) while others are focused on integrating data from multiple devices (NiMBaLWear [17], SKDH [18]). They are fully integrated packages, designed to calculate complex measures and summary reports using a range of technical approaches. Yet, they are all configured to output final digital measures rather than to assist the comparison and development of measures in a verified software environment which can satisfy the needs of clinical regulators. They do not support bout characterisation from raw data and can have opaque rules for managing imputation and interruptions. All have the common pre-processing elements of calibration, non-wear detection and basic measure characterisation which have been simplified into the *GENEAcore* package.

Returning to the first stage shown in Fig 1, the measurement period information contains all essential metadata including directions for correct file reading, unique data source identifiers to link records, calibration of the data to maximise sensitivity, determination of data completeness and the assessment of non-wear. The accurate determination of wear time is critical to understanding both participant adherence and data loss due to operational constraints [19]. A review of 14 non-wear detection algorithms has provided a consolidated approach with good accuracy and time resolution (S1 Table). In this study, we demonstrate the traceability of file handling, the performance of the non-wear detection approach and formally validate a widely used threshold for non-movement.

The second step in the pipeline is the decomposition of continuous time series data into a contiguous series of time windows. This splits the sensor-level data into manageable chunks for characterisation, analysis and visualisation. Fixed-duration epochs are traditionally used as they were simple to implement on early wearables and are convenient for researchers [20]. However, there is no intrinsic relationship between the events and the physical behaviours under investigation because the timing is arbitrary. The selection of the epoch duration is also difficult; too short and there is not enough data to characterise a behaviour effectively, too long and they contain multiple behaviours. Changes in estimates of time spent at various activity levels with different epoch lengths are well reported [21, 22] acknowledging interactions with activity level cutpoints [23] and the subsequent impacts on public health guideline adherence [24] and associations with related outcomes [20, 25]. Variable duration events [26] based on transitions [27] identified from the variability of the data itself gives an alternative approach. This method has the advantage of ensuring that different, consecutive behaviours are not summarised into a single time window - improving accuracy and reducing confusion in later classification stages.

The concept of a bout was originally used in public health guidelines based on a minimum duration of physical activity that was linked to health benefits [28]. Increasingly, bouts are used to analyse all types of physical behaviours (for example, movement patterns [29]) with variable minimum durations [30, 31] and different bout criteria [32]. When working with epochs, bouts need to be reconstructed with, often, complicated rules for epoch inclusion and allowances for interruptions. Events provide a more natural transformation to behavioural bouts but may identify more interruptions than long epochs depending on the minimum event duration specified. In this study, we determine optimal parameters for data-driven event detection and investigate the resulting physical behaviour event durations. We discuss how event approaches may help to resolve the challenges with epoch length selection and bout rules.

In the third step of Fig 1, the sensor-level measurements are processed to characterise the epoch or event. This involves the application of mathematical operations to each of the individual sensor measurements to describe the data in some way. Often, several features (ideally, independent) will be extracted for a multi-dimensional perspective. There are many standard, open algorithms that can be applied [33], and the pipeline flexibly supports the verified implementation of novel, raw data functions. In this study, we will show the utility of our framework by comparing two commonly used techniques for estimating activity intensity in both the epoch and event paradigms.

The pre-processing pipeline is designed to establish consistent inputs for the remaining 2 steps beyond the scope of this work. Digital measures may be based directly on the data characteristics at this stage (for example, mean activity intensity) or rely on further manipulation and modelling to infer physical behaviours that would be recognisable to an observer (for example, sleep or walking). Ultimately, these measures are often aggregated into summaries of daily living activities over multiple days to represent the lifestyle of the participant (for example, total active duration).

Without pre-processing as a distinct and independent layer, the validation of a digital measure becomes inseparable from its base assumptions and calculation processes. A digital measure then cannot be compared independently of the package that created it. The risk is that we become locked into a plurification of end-to-end processing pipelines, such as those mentioned earlier, for every digital measure or have different interpretations of an individual’s physical behaviours depending on the pre-processing used.

## Methods

The study used triaxial acceleration and temperature data from the GENEActiv Original (ActivInsights Ltd, Kimbolton, United Kingdom) in the raw output ‘.bin’ format that contains all measurements and recording information. These were processed in the open-source R package, *GENEAcore* (https://cran.r-project.org/package=GENEAcore), hosted by CRAN (The R Foundation, Vienna, Austria). The work was completed by a cross-functional team of academic, industry and specialist software personnel.

Engineering verification of the software was completed through unit and integration testing with design controls (as described in ISO62304[34]) and algorithm code inspection by relevant experts in the team. Development for the package was undertaken across Windows and macOS utilising R Studio and its associated R package development libraries *dev tools, usethis* and *testthat*. Source code control, package building and continuous integration testing were performed using GitHub and GitHub Actions. Unit testing utilised independently calculated values or reference results from previously published calculations and packages providing 90% code coverage by file. Builds were tested across a range of platforms including Windows, macOS and Linux. Package build and final release test approval is completed by automated build routines in clean, independent environments. The measure outputs were validated against the *GGIR* and *GENEAread* R packages (also hosted by CRAN) to ensure equivalence.

Verification testing was performed using multi-day, wrist-worn datasets that included 68 participants and 69 unique devices from 100 files. Analytical validation testing was completed with objective and observed criterion measure data for 36 participants from 65 files (Table1). The datasets were selected from available resources to include a wide range of ages and a reasonable gender balance.

Ethical approval for the HeLP study was obtained from the Peninsula College of Medicine and Dentistry in March 2012 (reference number 12/03/140). The Institutional Review Board and Regional Ethics Committees of Copenhagen University Hospital Rigshospitalet and Amsterdam University Medical Centre approved the SafeHeart study protocol. Permission has been granted by the first author for the use of personal data for this study. The polysomnography studies were approved by the University College London ethics committee and the NRES Committee North East Sunderland ethics committees. ActivInsights have authorised the use of the office-collected non-wear data and confirmed it was collected with consent under appropriate safeguards.

All data processing and statistical tests are processed in R version 4.4.2 [38].

### Measurement period information

The measurement period information is initially extracted from a bin file by the *GENEAcore* package, which then implements an optimised file read procedure to extract 1-second downsampled time series data for calibration, non-wear detection and transition detection.

#### Calibration

A rolling standard deviation is calculated from the 1-second downsampled accelerations over a defined window of 120 seconds, with a threshold of 13m*g* applied to the 3 axes mean identifying measurements of non-movement. These selected measurements are iteratively fitted to a sphere to estimate auto-calibration figures from real world data [39]. The values selected for all processing parameters are shown in S2 Table with references.

All multi-day files are auto-calibrated using this method with the number of iterations required and the improvements in calibration, as calculated by residual error, are recorded. The laboratory-generated multi-orientation datasets both with 53 periods of non-movement each of about 20 minutes over 12 hours are used to evaluate the correct convergence of the auto-calibration. Once the calibration was completed, the scale and offset elements are systematically perturbed and the residual error calculated (+/-1% scale and +/-50% offset).

#### Non-wear detection

Many approaches to the detection of non-wear have been published since the advent of raw data acceleration measurement and a summary of algorithms is shown in S1 Table. These are normally for single devices although dual device systems are known [40]. The methods use features derived from acceleration across 3 axes in varying combinations with temperature and, occasionally, time of day.

While signal properties can change between device and sensor models, the standard deviation of acceleration appears to offer a consistent level of detection performance robust to poor calibration [41]. Low pass signal filtering, including epochs and running means, allow absolute values of acceleration to be used [42, 43] and posture changes to be determined [44, 45]. However, using filtered signals in subsequent analysis reduces dynamic range [46]. Some publications note that the z axis has different noise characteristics and temperature dependencies compared to the other axes [47, 48]. This can be explained by the construction of the sensor with the z axis being through the plane of the silicon layers as opposed to in-plane for the x and y axes.

The temperature sensors within devices can be used with absolute thresholds [43, 47, 49, 50], but have complex signal characteristics over time. While the thermal lag due to the mass of the wearable would suggest a first order response, the initial rate of change is suppressed by the internal hysteresis of the sensor and quantization of signal output [51]. Publications report seeing maximum changes in temperature on device removal of 2-3 °C after 3-5 minutes [43, 52, 53] and a decoupling from body temperature after 16 minutes [49]. These findings help to define an optimal time window for detecting device removal using temperature changes.

Earlier algorithms all use fixed length epochs with varying overlaps but, more recently, event-based approaches have been deployed with transition detection of the acceleration [43, and temperature signals [52, 53]. Event-based approaches have the advantage of precise output time resolutions without needing to use very short epoch lengths which are prone to classification error [55].

Several algorithms use ‘boarding rules’ to update the assessment of non-wear events and neighbouring events once an initial classification has been made. These can be based on the duration of the non-wear events [44, 56], the duration of the gaps between non-wear events [54] or the signal characteristics of the event [57]. In addition, these approaches can be combined with sleep determination [44, 57], recognising this as the most likely behavioural confusion. Different rule sets are applied during night periods or when proximal events are determined to be sleep.

The increased performance levels of newer approaches (resolution, accuracy and minimum detection periods) have come at the expense of processing complexity. The *GENEAcore* non-wear algorithm aims to simplify and reduce processing times without compromising accuracy. Importantly for interoperability and modularity, the detected non-wear periods are stored in the metadata separately from the time series measurements. It uses approaches, features and parameter values that have been successfully and consistently used in the prior art described above (S1 Table and S2 Table). It is an event-based approach with candidate events identified when the previously derived, 2-minute rolling average of 1-second downsampled acceleration standard deviation drops below 13m*g* [58]. Boarding rules are used on candidate events to combine events that are separated by less than 5 minutes. These events are classified as non-wear when any of these rules are satisfied:

- Temperature drop due to device removal – more than 0.7 °C per minute drop in the first 4 minutes of non-movement with less than 2 posture changes; or
- Sustained non-movement without posture change – any single event that is greater than 2 hours in duration; or
- Long-running non-movement – continuous non-movement periods greater than 12 hours in duration irrespective of the number of posture changes.

The resulting outputs have a minimum detectable non-wear event duration of 5 minutes (timespan of temperature drop rule plus rolling average lag) with a time resolution of 1 second. These are stored in the measurement period information.

Here, we test the non-wear algorithm with wrist-worn data against both the observer diaries of controlled non-wear and sleep clinic participation. The office non-wear and sleep polysomnography criterion data are processed to create estimates for non-wear for each second. The estimates are then matched with observer records to ascertain true positive, true negative, false positive and false negative rates which are combined into balanced accuracy. A sensitivity analysis investigated the impact of changes in the acceleration standard deviation threshold on the classification accuracy.

### Epochs and events

Epochs and events are specified by their integer second start time and duration. Epochs start from the beginning of the first minute of the day. The transition detection for event start times utilises the 1-second downsampled time series from the measurement period information and the resulting transition time points are stored there.

#### Transition detection

*GENEAcore* uses Pruned Exact Linear Time (PELT) changepoint detection [59] from the *Changepoint* R package to identify transitions in the mean and variance of downsampled acceleration. Each axis is assessed separately with the resulting timestamps combined while amalgamating transitions within 5 seconds of each other (minimum event duration).

The PELT method uses a penalty value to control sensitivity to changes in the data and, therefore, the average number of transitions detected. Accelerometer studies in animals have previously used sample entropy to select penalty values that maximise the information density of the event summary [60]. Here, we calculated the root mean square error (RMSE) of the event summary output compared to the 1-second epoch output on each axis for different penalty values on a day randomly selected from each of the 100 verification files. Inflection points in the relationship between the average number of events per day and RMSE represent the point where a stable representation of the original signal has been reached.

Once suitable penalty values are selected for each axis, the 2 multi-orientation laboratory datasets are used to test the method. The timings of true transitions are compared to those determined by the PELT algorithm. The impact of varying the minimum event duration on the average number of events per day are then calculated. Additionally, we investigate the distribution of event durations.

### Characterisation

At this point in the processing, the measurement period information contains sensor calibration parameters, non-wear periods and event transitions. The raw, sensor-level measurements can now be processed as events or epochs with the selected algorithms to characterise the data. Any further processing to classify the events or epochs (for example, as sleep or high intensity activity) is a further step beyond the scope of pre-processing (Fig 1).

The architecture of the *GENEAcore* package allows different methods to be executed within a consistent processing environment. Here, we compare two activity intensity measures and different approaches to aggregation.

The earliest raw acceleration summary algorithm [61] took the sum of the gravity-subtracted vector magnitudes in an epoch (SVM_gs_) using the absolute value of any negative numbers. The resulting measure can be difficult to interpret as it is linearly dependent on both the sample frequency and epoch length; the mean of the absolute gravity subtracted acceleration (AGSA) addresses this drawback. Euclidean norm minus one (ENMO) was later proposed [56] and differs only by setting negative numbers from the subtraction of gravity to zero. Both AGSA and ENMO are commonly expressed in milli-units of gravity, m*g*, where *g* is equal to 9.81 m/s^2^.

AGSA and ENMO are calculated for all time points in the verification dataset using 1-second epochs as this is recognised to give the most accurate assessment of instantaneous intensity [62, 63]. The percentage overlap of each above and below the 13m*g* standard deviation movement threshold were calculated to understand their ability to discriminate effectively in low movement conditions. AGSA and ENMO are then aggregated by day to investigate the concordance [64, 65] of assessed active duration using a previously defined AGSA threshold of 62.5m*g* [61, 66] and the modelled ENMO equivalent.

Despite the absence of an intensity criterion measure, it is still possible to illustrate how an independent pre-processing stage supports future research. Event outputs are also calculated for each day in the verification dataset as well as the epoch outputs. Both are then aggregated by day to investigate the differences between variable duration event and fixed duration 1-second epochs aggregation on estimates of daily active duration.

## Results

The *GENEAcore* pre-processing pipeline was configured to output 3 data objects: the measurement period information, 1-second epochs and variable duration events. Once completed, the measurement period information contained calibration values, non-wear periods and detected transitions separated from both the original data source and output time-series data. These objects were traceably linked by a unique source identifier derived from source file itself, rather than any manually controlled file name.

### Calibration

During auto-calibration of the verification datasets, the mean number of iterations required to meet the incremental fit improvement threshold (see S2 Table) was 51.3 (31.6 SD). The mean improvement from factory calibration was 9.5m*g* resulting in a mean residual sphere fit error of 6.7m*g* (0.48m*g* SD).

All calibration perturbations of the laboratory-generated multi-orientation datasets resulted in an increased residual sphere fit error, confirming convergence. The mean increase for a 1% change in scale was 1.1m*g*, and 1.2m*g* for a 50% change in offset.

The repeatability of the auto-calibration was evaluated on the 2 devices worn multiple times by the lead author. The mean standard deviation of calibration gain and offset between devices was 19.5m*g* and 31.5m*g*, while the within device mean standard deviation was 1.0m*g* and 2.1m*g*, respectively.

### Non-wear detection

The sleep polysomnography (PSG) and office non-wear criterion data were processed to create estimates for non-wear for each second. The office non-wear contained 157 observed non-wear events with a mean duration of 1268 seconds (min. 178, max. 3629). All assessments of non-wear during polysomnography measurements were assumed to be false positives. Table 2 shows the resulting sensitivity, specificity and balanced accuracy alongside the base observation rates as described in the methods.

**Table 1.** Verification and validation datasets.

| Dataset | Measurement period & data | Number of files | Mean age | Male % |
| --- | --- | --- | --- | --- |
| **Verification testing** |  |  |  |  |
| Randomly selected participants from the HeLP study [35] | 9 days @ 85.7Hz | 33 | 9.75 yrs | 49% |
| Randomly selected participants from the SafeHeart study [36] | 17 days @ 50Hz | 17 | 64 yrs | 81% |
|  | 30 days @ 20Hz | 17 |  |  |
| Sequential recording periods from the lead author of this study | 15 days @ 100Hz | 33 | 49.5 yrs | 100% |
| **Analytical validation** |  |  |  |  |
| Laboratory-generated multi-orientation datasets | 12 hours @ 100Hz | 1 | n/a | n/a |
|  | 12 hours @ 10Hz | 1 |  |  |
| Nights with simultaneous sleep polysomnography [37] | 1 night @ 85.7Hz | 55 | 44.9 yrs | 61% |
| Office-collected non-wear following an observer-controlled protocol | 8 hours @ 100Hz | 8 | 33.9 yrs | 25% |

Descriptions of the data and participants used in the verification testing and analytical validation.

**Table 2.** Non-wear detection performance.

|  | True positives | True negatives | False positives | False negatives | Sensitivity | Specificity | Balanced accuracy |
| --- | --- | --- | --- | --- | --- | --- | --- |
| Sleep PSG | 0 | 1.8x10<sup>6</sup> | 906 | 0 | — | 99.95% | — |
| Office non-wear | 66564 | 118240 | 2326 | 11918 | 84.8% | 98.1% | 91.4% |
| Combined | 66564 | 2.0x10<sup>6</sup> | 3232 | 11918 | 84.8% | 99.8% | 92.3% |

Results of non-wear detection algorithm testing in different wear scenarios.

The acceleration standard deviation threshold of value was varied between 6mg and 20mg to test whether the widely used 13mg threshold [58] for identifying non-movement was optimal for this sensor and non-wear algorithm in these wear scenarios. The changes in detection performance are shown in Fig 2, where a marginal increase in balanced accuracy from 92.3% to 92.6% can be seen at an acceleration standard deviation threshold of 12m*g*.

**Fig 2.**
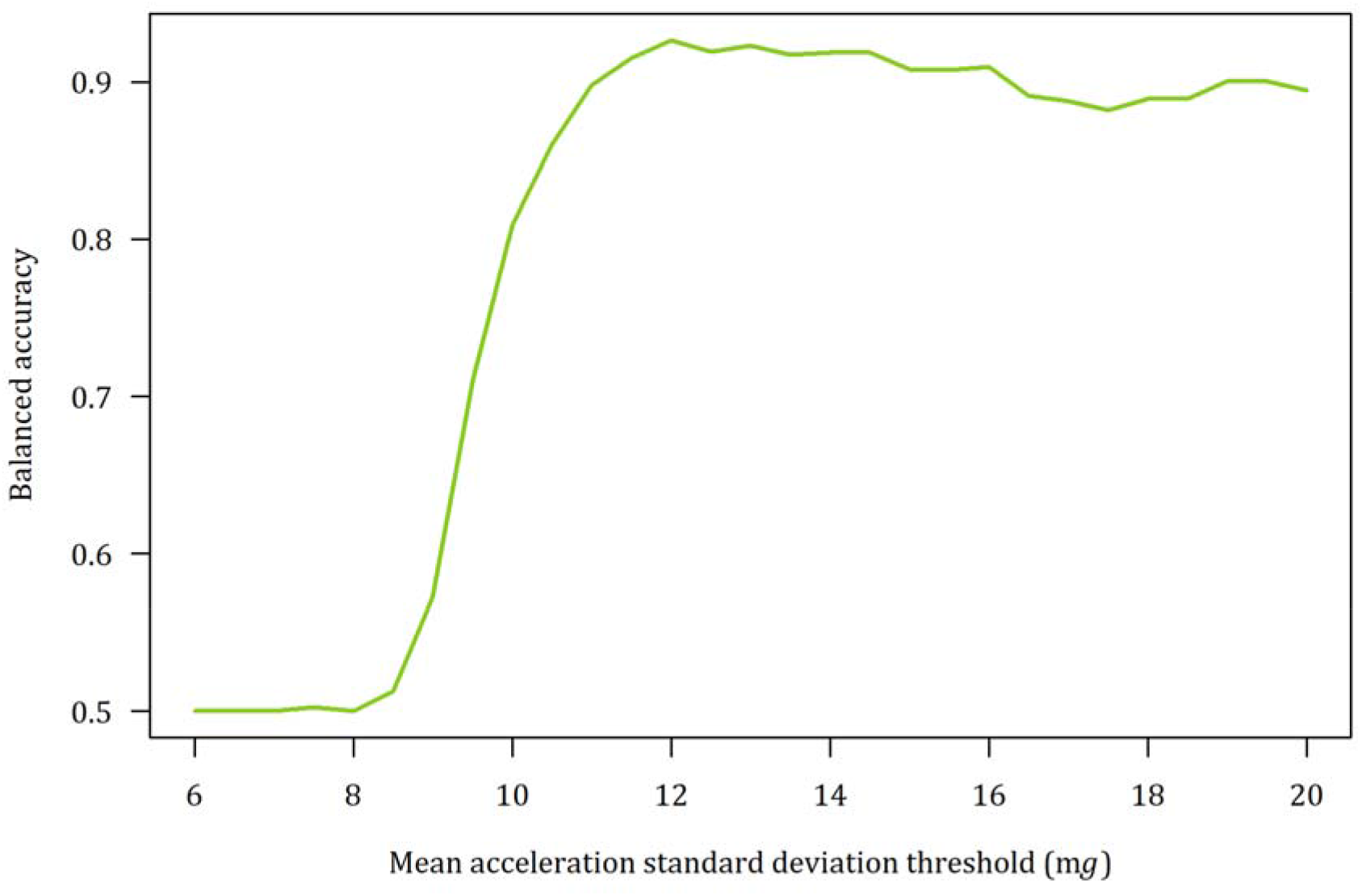
Non-wear detection sensitivity analysis. Changes in non-wear detection performance as measured with balanced accuracy for different values of mean acceleration standard deviation threshold.

### Transition detection

Optimal penalty values for PELT changepoint detection were found to be 18, 25 and 16 for the x, y and z axes respectively. When validated against the 2 multi-orientation datasets, only 1 of 104 transitions were not correctly identified and mean minimum distance to transitions was 1.65 seconds.

For each axis individually, the target number of events per day, based on the RMSE inflection points, was approximately 500. When the detected transitions for each axis in the verification dataset were combined with a minimum event duration of 5 seconds, the mean number of events per complete day (n=1,296) was 1,116 (221 SD). S1 Fig shows plots of RMSE, events per day and penalty value with inflection points for each axis and verification dataset sub-group.

The minimum permitted event duration when combining transitions was varied from 3 to 60 seconds with plots of resulting transitions per day for each verification dataset sub-group shown in Fig 3.

**Fig 3.**
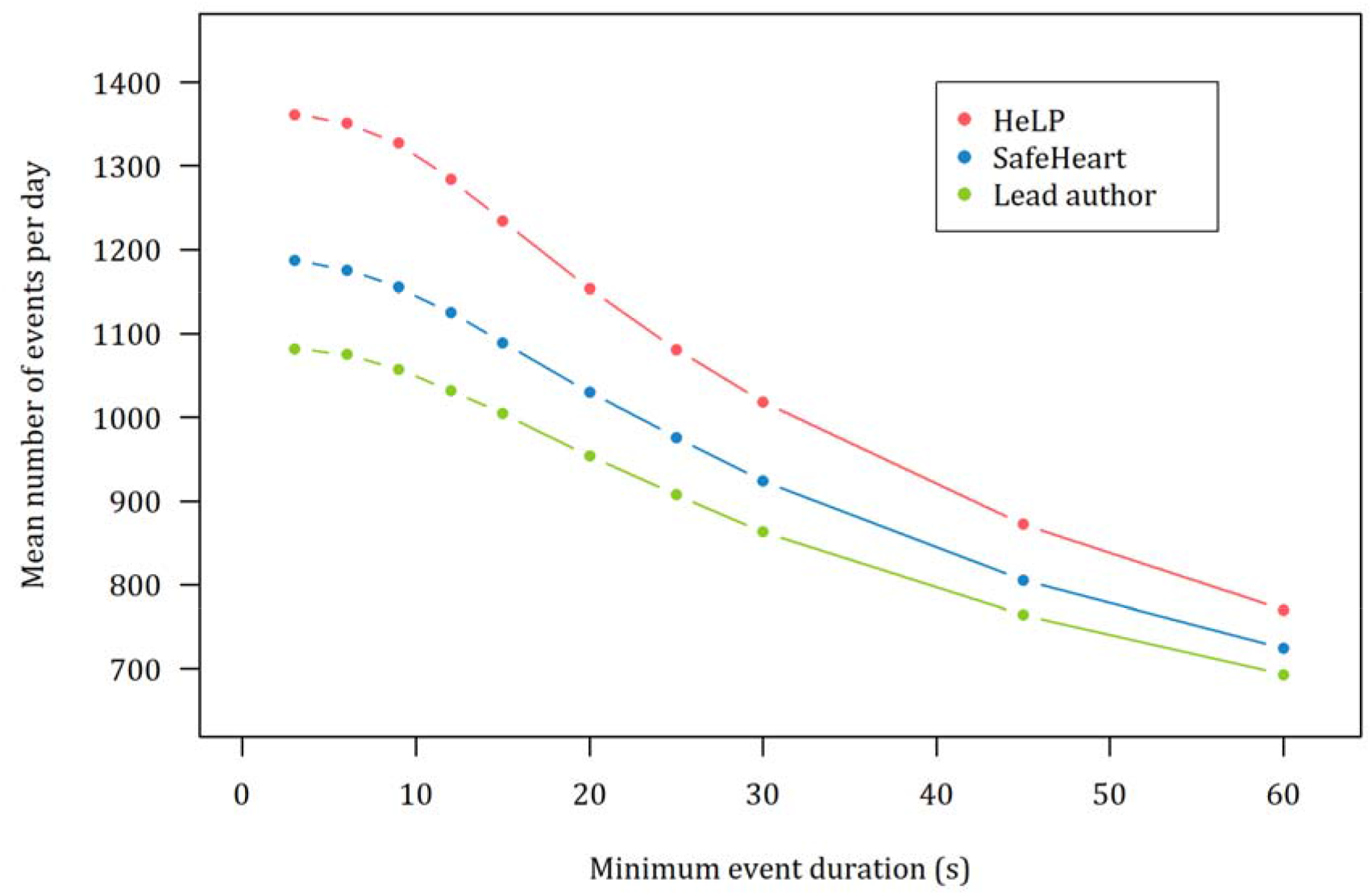
Impact of minimum event duration on events per day. The variation of the number of daily events with minimum event duration in the 3 verification population groups.

The distribution of event durations approximated closely to the log normal (μ = 3.9, σ=1.1) giving an expected event duration, E(X), of 68.6 seconds.

Data summaries based on variable length events provided a compression ratio of 87.9 compared to a 1-second fixed epoch output, while preserving a 1 second resolution for changes in the underlying data.

### Characterisation

The relationship between AGSA and ENMO is shown in Fig 4 for every second in the verification dataset (130,484,716 data points). The red group has all time classed as moving (mean standard deviation of acceleration above 13m*g*) and the blue group contains the remaining time classed as non-movement. Above 1g of mean acceleration, AGSA and ENMO give near identical results. In the lower acceleration ranges of the moving group, ENMO consistently under-reports compared to AGSA. In the non-movement group, the asymmetric treatment of negative values in the ENMO algorithm is apparent, with values of mean acceleration often reported below 1m*g* despite the 6-7m*g* noise floor of the sensor. The overlap of AGSA values in the moving and non-movement groups was 6.0% while the ENMO overlap was 10.5%.

**Fig 4.**
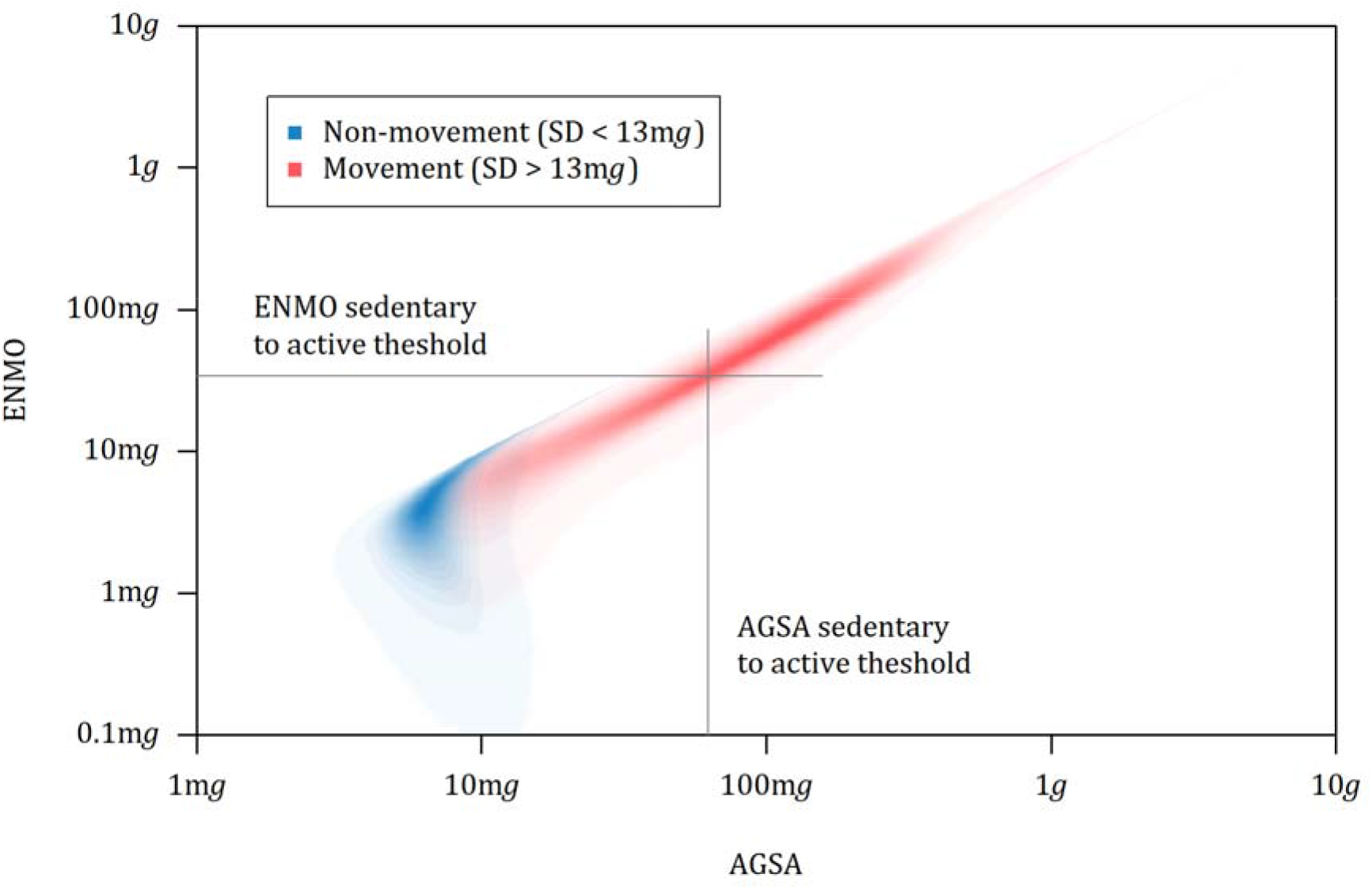
Comparison of activity intensity calculated by AGSA and ENMO. Density plot of 1s epochs summarised by AGSA and ENMO for the verification dataset split into groups of non-movement (blue) and movement (red).

The typical sedentary to active threshold for AGSA of 62.5m*g* [61, 66] is shown vertically in Fig 4. A linear regression of the movement group (r^2^ = 0.957, slope = 0.872, intercept = -20.5 m*g*) establishes an equivalent threshold for ENMO of 34.1mg, shown horizontally.

When these sedentary to active thresholds are applied daily over 1,296 complete days, the concordance of the resulting daily active duration was found to be greater than 0.99. Correlation and Bland-Altman plots are shown in S2 Fig. ENMO has bias of +1.25 minutes per day compared to AGSA.

The *GENEAcore* pre-processing pipeline allows us to contrast event and epoch segmentation. The correlation of active duration per day determined with variable duration events and fixed duration 1-second epochs is shown in S3 Fig with an accompanying Bland-Altman regression plot. The concordance was 0.831 (95% CI, 0.821–0.842), however the Pearson correlation coefficient was 0.985 (95% CI, 0.983–0.986). The regression model of the differences shows the event-derived active durations per day to be consistently 31.5% (95% CI, 30.8 - 32.2) higher than those from 1-second epochs.

## Discussion

Our aim in this study was to define and test a simplified, modular pipeline for the pre-processing of high-resolution accelerometry – assuring the quality of raw, sensor-level data as they are successively refined into digital measures.

The handling of data objects during the study demonstrated full traceability from input data sources to the output digital measures with a record of all processing steps in the measurement period information. The modularity of implementation and reporting provides the required transparency, while also supporting easy application of new or improved techniques. Integrated testing and change control will assure regulators of the suitability for health applications.

Results recorded for the calibration stage confirm the success of earlier studies [39], showing residual error comparable to the noise floor of the sensor itself. Furthermore, the perturbation tests demonstrate correct convergence of the optimised method and repeatability tests confirm that the calibration of these sensors is static over time. The detection of transitions using changepoint analysis proved reliable with 99% of transitions detected, on average, within 2 seconds of their occurrence.

This is the first study to empirically test the 13mg acceleration standard deviation threshold commonly used in the literature since 2014 [58]. This was achieved by calculating the balanced accuracy of non-wear detection through a range of threshold values. The results indicate a slight reduction of the threshold could be beneficial for this specific device, however preserving current practice affords some leeway for noisier sensors, with a minimal reduction in efficacy.

The non-wear algorithm components, all based on existing know-how, are independent of body location and show no variation between adults and children or male / female participants [49]. We consider that the approaches and results have high generalisability to wear position and population. The criterion datasets used to test the non-wear algorithm were selected to challenge in situations where performance can be marginal: sleep measured by PSG and short periods in a lived environment controlled by an observer. Of the previous non-wear validation studies, only 2 used observers [44, 57] and 1 used a separate device [54], all others relied on participant diaries or expert rating of data. The high overall specificity is excellent for sleep measurement where false positives for non-wear during the night are problematic for clinicians. The sensitivity to short off-wrist periods is reasonable; it will be higher and comparable to the other published methods for longer periods of non-wear where the participant is not in proximity with the device. The detection of non-wear during transport will remain difficult, however this is normally accompanied by long, still periods where accuracy is high.

The detailed comparison of 2 commonly used measures of activity intensity (AGSA and ENMO) reveal that attention must be given through every stage of digital measure selection and embodiment for any health application. A small change in the algorithm has major consequences. The sedentary AGSA threshold of 62.5m*g* [61, 66] was translated to an equivalent ENMO threshold of 34.1m*g*; a value which is exactly in line with other studies across a range of populations [67, 68]. Therefore, for higher movement levels above the sedentary thresholds, AGSA and ENMO are substitutable for measuring daily active times using the linear regression described. More care is required at lower movement levels, when analysing individual events and when using activity intensity as a continuous measure. Below the sedentary threshold, ENMO quickly becomes non-linear with reduced movement, outputting results inconsistent with the measurement noise of the sensor. There is more headroom for AGSA below the sedentary threshold until sensitivity is lost in a more linear fashion.

A particular contribution of this work is the comparison of epoch and event pre-processing made possible by the *GENEAcore* pipeline. Variable duration events created higher assessments of daily activity duration compared to those calculated from 1-second epochs. The disparity in outputs was an extra 19 minutes for every additional hour of activity. The magnitude of this disparity will vary with the epoch length selected for comparison. The threshold used in this analysis to identify active behaviours encompasses light, moderate and vigorous intensity physical activity. Longer epochs are consistently reported to reduce estimates of both vigorous and sedentary time [22, 23, 24, 25, 69], pushing all estimates towards the mid-intensity bands. So, the disparities in reported active time compared to events will decrease with longer epochs. However, in the epoch paradigm, these longer epochs will under-report vigorous active times, whereas variable duration events will still capture short bursts of higher intensity physical activity. This ability for event-based processing to adapt to the latent structures of the data provides a solution to suggestions that different epoch lengths are more effective at detecting behaviours of different intensity [20, 21].

Data-driven estimates of the typical durations of human movement behaviours are not widely available. The log normal distribution found here, with an expected event duration of just over a minute, is similar to that of other land-based mammals [60]. A minimum event duration of 5 seconds preserved the complexity of daily data, allows short bouts of activity to be recorded [69, 70] and gives a finite time for the transitions themselves [27]. These findings confirm the existing conventions of epoch length selection of between 1 and 60 seconds. The compromises associated with epoch length choices outlined above are resolved by variable duration encoding. The event-based paradigm delivers the timestamp accuracy of short epochs and efficient data compression of longer epochs with high classification confidence. In addition, the relative timings of the transitions themselves expose underlying patterns of physical behaviours.

The uncertainties associated with the selection of appropriate epoch length are compounded once epochs are grouped to represent bouts of physical behaviour. The *GENEAcore* pre-processing pipeline specifies event transitions prior to characterisation and classification – allowing an explicit translation from data-driven events to bouts without further processing. In contrast, existing epoch-only pipelines must apply additional rules with parameters to identify bout transitions, epoch aggregation and permissible interruptions. These rules require substantial verification and analytical validation for them to be used repeatably and with confidence.

At first glance, event approaches appear more complicated than the use of traditional epochs. The variable durations pose methodological problems in parameter selection, visualisation, aggregation and bridging of reporting boundaries (for example, across days). However, these obstacles are inherent to bout representations and data-driven event transition detection simplifies the later stages of process validation. Lastly, events defined in the early steps of pre-processing supply event duration as a valuable, additional feature for later classification stages.

Our study has limitations in the small sizes of datasets used for the analytical validation steps, absence of processing time comparisons between existing packages and the lack of a criterion measure for activity intensity. Future work can use other larger, previously collected data to further investigate comparative accuracies of AGSA vs. ENMO activity intensities and epoch vs. event pre-processing. Ideally, this work would use continuous measures of activity intensity rather than cutpoint-derived energy expenditure classes to avoid threshold effects. A better understanding of the impact of the rules for bout creation from epochs on the reporting of physical activity acquisition (durations, volumes and patterns) at different epoch lengths is needed. Data-driven events would provide a stable benchmark for this analysis. Study strengths include the use of a cross-functional team to create a maintainable, engineered tool with backward compatibility for existing data. Deliberately constraining scope to the pre-processing pipeline alone enhanced the depth and focus of results. The thorough documentation of assumptions will support the consistent implementation of existing measures identified as important to public health [71, 72] as well as clinical research.

## Conclusions

Our focus on verification and analytical validation of accelerometer pre-processing has exposed detailed engineering considerations that must underpin reliable digital measures. The rush towards results with health associations often leaves little room for these important topics.

The meticulous evaluation of the simplest steps of pre-processing has confirmed the attention to detail that is required to create robust and reproducible results in later clinical validation. The integrity of data pathways is preserved through standardised approaches and rigorous implementation.

We have shown this optimised framework to be a capable building block of future research, clinical care and public digital health systems. It can deliver the basic digital measures needed to understand physical behaviours over multiple days with minimum assumptions.

## Acknowledgements

The authors acknowledge the contributions of the study leaders and participants that provided data for the verification and analytical validation datasets, particularly: Katrina Wyatt and Jennifer Lloyd of the HeLP study; Diana My Frodi and Maarten Kolk of the SafeHeart study; and Vincent van Hees, Sarah Charman and Kirstie Anderson for the Newcastle polysomnography and accelerometer data. The authors would also like to thank Brad Metcalf of University of Exeter for his statistical support. For the purpose of open access, the authors have applied a Creative Commons Attribution (CC BY) licence to any Author Accepted Manuscript version arising from this submission.

## Supporting information

**S1 Table. Non-wear detection methods**. Descriptions of different methods for the detection of non-wear including: the general method, features used for detection, the minimum detectable non-wear duration, the time resolution of non-wear periods, the criterion used in their validation and the wear location.

**S2 Table. Processing parameters**. The definitions, values and the supporting references for the parameters used in the pre-processing operations.

**S1 Fig. Changepoint penalty value assessments**. The variation of the number of daily events and the RMSE of intensity with changepoint penalty value for each axis of acceleration in the 3 verification population groups.

**S2 Fig. Comparing daily active duration using AGSA and ENMO**. (a) Correlation plot of active hours per day as assessed by AGSA and ENMO, (b) Bland-Altman plot of the means and differences of AGSA and ENMO active hours per day with mean bias and 95% SD confidence intervals.

**S3 Fig. Comparing daily active duration using event and epoch aggregates of AGSA**. (a) Correlation plot of active hours per day as assessed by AGSA 1s epoch and variable duration events, (b) Bland-Altman regression plot modelling the differences of AGSA 1s epoch and variable duration event active hours by event-assessed duration per day with mean bias and 95% SEE confidence intervals.

