## Supplementary Figure 1 for "Validation and optimisation of wearable accelerometer data pre-processing for digital measure implementation and development"

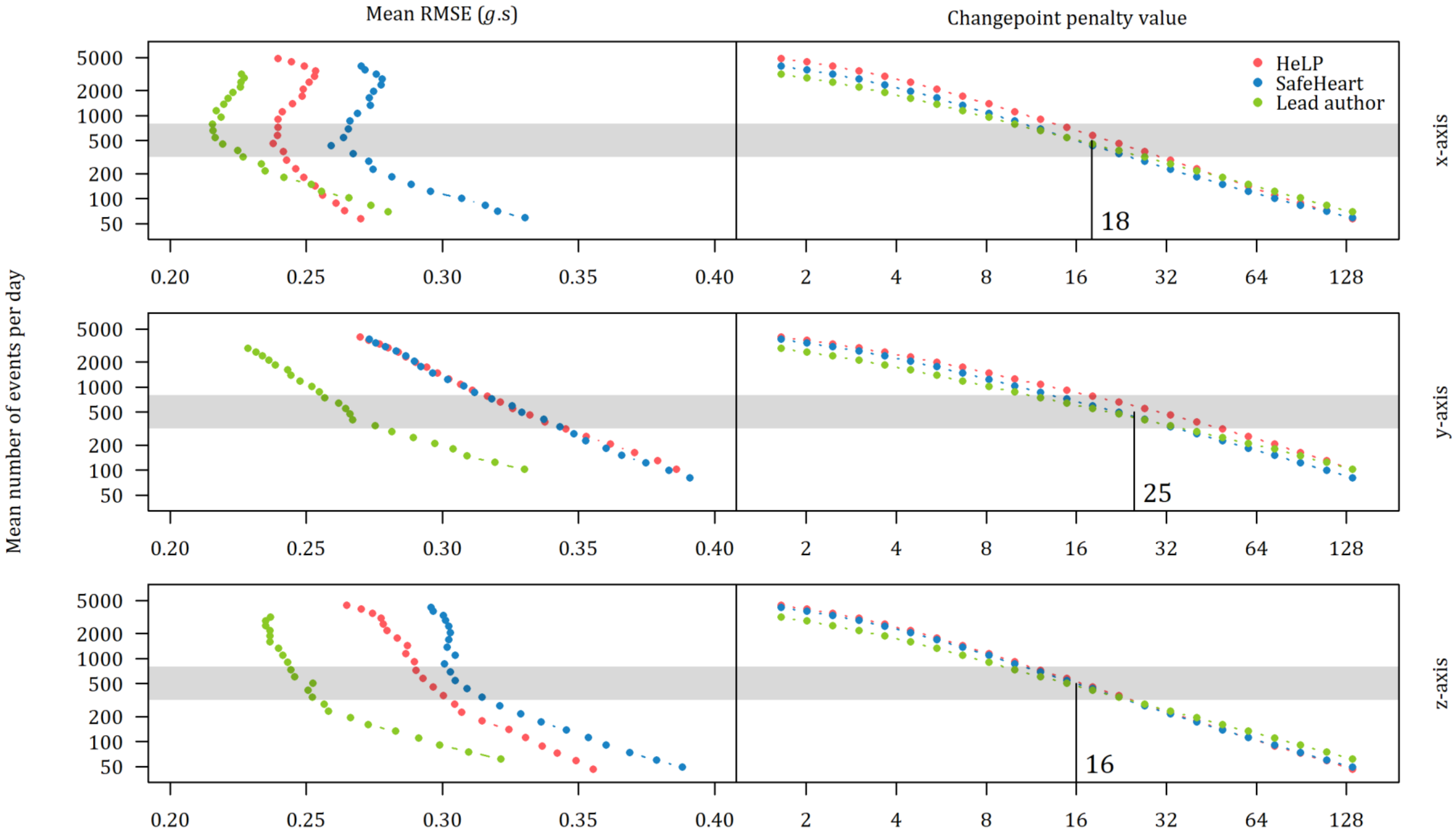


**S1 Fig. Changepoint penalty value assessments.** The variation of the number of daily events and the RMSE of intensity with changepoint penalty value for each axis of acceleration in the 3 verification population groups.
