## Supplementary Table 1 for "Validation and optimisation of wearable accelerometer data pre-processing for digital measure implementation and development"

**S1 Table. Non-wear detection methods.**

| Authors | Method description | | | | |
|---|---|---|---|---|---|
| | Features | Minimum detection | Resolution | Criterion measure | Wear location |
| van Hees et al. 2011 [41] | Consecutive epochs of 30 minutes are used to calculate the mean and range of each of the acceleration axes. An epoch is classified as non-wear if the standard deviation is less than 3m*g* or range is less than 50m*g*, for at least two out of three axes. | | | | |
| | Acceleration SD & range | 30 minutes | 30 minutes | - | Wrist |
| van Hees et al. 2013 [56] | Epochs of 60 minutes with a 75% overlap are classified as non-wear if the standard deviation is less than 3m*g* or range is less than 50m*g*, for at least two out of three axes. Bordering rules are iteratively applied to non-wear periods of 3 hours and 6 hours based on post and prior non-wear. | | | | |
| | Acceleration SD & range | 60 minutes | 15 minutes | - | Wrist |
| da Silva et al. 2014 [58] | Epochs of 60 minutes with a 75% overlap are classified as non-wear if the standard deviation is less than 13m*g* or range is less than 50m*g*, for at least two out of three axes. | | | | |
| | Acceleration SD & range | 60 minutes | 15 minutes | - | Wrist |
| Zhou et al. 2015 [49] | Epochs of 1 minute incremented by 1 second are classified as non-wear if the temperature is below 26°C and acceleration standard deviation is less than 13m*g* in all three axes. | | | | |
| | Acceleration SD, absolute temperature | 15 minutes | 1 minute | Participant diary | Wrist |
| Ahmadi et al. 2020 [44] | Epochs of 30 minutes incremented by 1 second are classified as non-wear using different rules: standard deviation less than 13m*g*, high-pass filtered sum vector magnitude greater than zero and posture change. Bordering rules are applied to non-wear periods of 30 minutes or less. | | | | |
| | Acceleration SD, high-pass filtered SVM, posture changes | 10 minutes | 30 seconds | Observer diary | Wrist |

| | | | | | |
|---|---|---|---|---|---|
| Rasmussen et al. 2020 [50] | Epochs of 10s where the standard deviation of acceleration is below 20m*g* are combined and classified as non-wear if they are longer than 120 minutes, or 45-120 minutes with an average temperature below a threshold, or 10-45 minutes with an average temperature below a threshold and an end time in the expected wake time. | | | | |
| | Acceleration SD, time of day, temperature | 10 minutes | 10 seconds | - | Hip & thigh |
| Barouni et al. 2020 [57] | Epochs of 10 minutes incremented by 30 seconds are classified as non-wear if the power of movement frequencies between 0.1Hz and 0.4Hz is less than 2 x $10^{-5}$. Periods of continuous non-wear are checked to ensure they meet the above criteria at least 75% of the time. | | | | |
| | Acceleration spectral power | 10 minutes | 30 seconds | Observer diary & expert rating | Wrist & chest |
| Rahimi-Eichi et al. 2021 [42] | Epochs of 1 minute are used in a rolling 150-minute window. The standard deviation of each axis in each epoch is calculated. Periods are defined as non-wear when the root mean square of the moving average is below 18.5m*g* for all axes. | | | | |
| | Acceleration SD | ~ 150 minutes | 1 minute | Participant diary | Wrist |
| Sundararajan et al. 2021 [45] | Epochs of 30s are used to calculate features of posture, posture change, mean acceleration and 30 minutes smooth activity score. A random forest machine approach is used to determine non-wear with posture, posture change in bordering epochs and activity scores being the most important features. | | | | |
| | Posture, posture changes, activity score | 5 minutes | 30 seconds | Participant diary & expert rating | Wrist |
| Syed et al. 2021 [54] | Candidate non-wear events are created from epochs of 1 minute incremented by 1 second where the standard deviation of acceleration is below 4m*g*. Bordering non-wear events within 5 minutes of each other are merged. Raw acceleration data for the 3 second periods bordering the candidate non-wear event are processed by a convolutional neural network to detect wear start and wear end. | | | | |
| | Raw acceleration in a convolutional neural network | 1 minute | 1 second | Separate wearable device | Hip |
| Pagnamenta et al. 2022 [52] | Events starting and ending with a temperature change of 3°C over 5 minutes are classified as non-wear if the standard deviation of the band-pass filtered sum vector magnitude is less than 13m*g*. | | | | |
| | Acceleration SD, rate of change of temperature | 5 minutes | 1 minute | Participant diary | Lower back |

<table>
<tr><td rowspan="2">Vert et al. 2022 [43]</td><td colspan="5">Epochs of 5 minutes incremented by 1 minute using a low-pass filtered temperature signal and a 4-second rolling standard deviation of acceleration. Non-wear starts when the 1-minute standard deviation of acceleration is less than 8mg for 2 or more axes, standard deviation of acceleration is less than 8mg for 2 or more axes 90% of the time over 5 minutes, temperature is dropping more than 0.2 °C per minute and temperature is below 26 °C. Non-wear ends when the 1-minute standard deviation of acceleration is greater than 8mg for all axes, standard deviation of acceleration is greater than 8mg for 2 or more axes for 50% of the time over 5 minutes, temperature is rising more than 0.1 °C per minute and temperature is above 26 °C.</td></tr>
<tr><td>Acceleration SD, temperature, rate of change of temperature</td><td>5 minutes</td><td>1 minute</td><td>Expert rating</td><td>Wrist</td></tr>
<tr><td rowspan="2">Skovgaard et al. 2023 [47]</td><td colspan="5">Epochs of 10 seconds are used to calculate features of standard deviation of acceleration, mean acceleration, temperature and time of day. These are then processed by a machine learning decision tree to classify non-wear.</td></tr>
<tr><td>Acceleration SD & mean, temperature, time of day</td><td>10 seconds</td><td>10 seconds</td><td>Expert rating</td><td>Hip, thigh & wrist</td></tr>
<tr><td rowspan="2">Barakat et al. 2024 [53]</td><td colspan="5">1-minute epoch summaries of acceleration and temperatures are used to create a rolling 3-minute average temperature change and rolling 2-minute averages of acceleration standard deviation. Non-wear events are initiated by a concurrent −0.2°C drop in average temperature and an average acceleration standard deviation below 13 mg across all 3 axes. Non-wear events are terminated by an average temperature increase of 0.1 °C accompanied by an increase of the acceleration standard deviation of any of the three axes above 13 mg. Only validated during day wear.</td></tr>
<tr><td>Acceleration SD, rate of change of temperature</td><td>3 minutes</td><td>2 minutes</td><td>Participant diary</td><td>Arm sling</td></tr>
</table>

Descriptions of different methods for the detection of non-wear including: the general method, features used for detection, the minimum detectable non-wear duration, the time resolution of non-wear periods, the criterion used in their validation and the wear location.
