## Supplementary Figure 2 for "Validation and optimisation of wearable accelerometer data pre-processing for digital measure implementation and development"

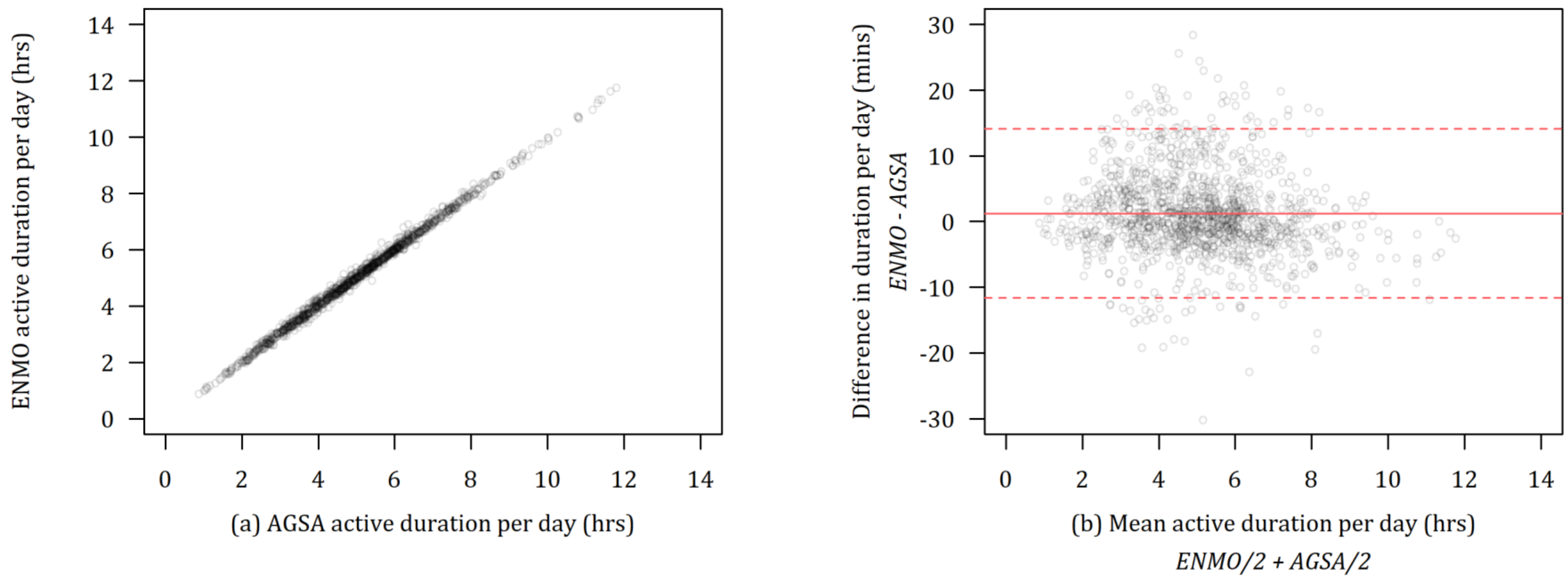


**S2 Fig. Comparing daily active duration using AGSA and ENMO.** (a) Correlation plot of active hours per day as assessed by AGSA and ENMO, (b) Bland-Altman plot of the means and differences of AGSA and ENMO active hours per day with mean bias and 95% SD confidence intervals.
