## Supplementary Table 2 for "Validation and optimisation of wearable accelerometer data pre-processing for digital measure implementation and development"

**S2 Table. Processing parameters.**

| Identifier | Units | Value | Technical description | References |
|---|---|---|---|---|
| still_seconds | s | 120 | The number of seconds included in the rolling standard deviation calculation of non-movement from 1Hz downsampled data. | Barakat et al. 2024 [53] |
| sd_threshold | *g* | 0.013 | The threshold applied to the rolling standard deviation of mean acceleration standard deviation of 1Hz downsampled data to determine non-movement. | da Silva et al. 2014 [58] with formal validation in this work. |
| spherecrit | *g* | 0.3 | The minimum required acceleration value for each axis in both directions to ensure sufficient range of non-movement positions for auto-calibration to be reliable. | van Hees et al. 2014 [39] |
| maxiter | count | 500 | The maximum number of sphere fit iterations attempted during auto-calibration to converge. | Originally 1000 in van Hees et al. 2014 [39] and optimised in this work. |
| tol | *g* | $1 \times 10^{-13}$ | The limit of incremental sphere fit improvements before auto-calibration is considered complete. | Originally $1 \times 10^{-12}$ in van Hees et al. 2014 [39] and optimised in this work. |
| temp_seconds | s | 240 | The number of seconds included in the rolling temperature difference calculation for non-wear, which also determines the shortest detection duration. | Vert et al. 2022 [43], Pagnamenta et al. 2022 [52], Barakat et al. 2024 [53] |
| delta_temp_threshold | °C | -0.7 | The threshold applied to the rolling temperature difference to determine non-wear. | Vert et al. 2022 [43], Pagnamenta et al. 2022 [52], Barakat et al. 2024 [53] |
| x_cpt_penalty | - | 18 | The manual penalty value applied in the PELT changepoint algorithm for the x axis. | Defined and formally validated in this work. |
| y_cpt_penalty | - | 25 | The manual penalty value applied in the PELT changepoint algorithm for the y axis. | Defined and formally validated in this work. |
| z_cpt_penalty | - | 16 | The manual penalty value applied in the PELT changepoint algorithm for the z axis. | Defined and formally validated in this work. |
| minimum_event_duration | s | 5 | The minimum interval between changepoint transitions. | Twaites et al. 2020 [27] and optimised in this work. |
| border_seconds | s | 300 | The maximum number of seconds between non-movement events for them to be combined into the same period. | van Hees et al. 2011 [41], Ahmadi et al. 2020 [44] and optimised in this work. |

| | | | | |
|---|---|---|---|---|
| long_still_seconds | s | 7200 | The number of seconds for any single non-movement event beyond which the whole period is classed as non-wear. | Rahimi-Eichi et al. 2021 [42] and optimised in this work. |
| posture_changes_max | count | 2 | The maximum number of adjoining non-movement events that make up a single period of non-wear less than the maximum non-move duration. | Ahmadi et al. 2020 [44], Sundararajan et al. 2021 [45] and optimised in this work. |
| non_move_duration_max | s | 43200 | The number of seconds beyond which non-movement periods are automatically classed as non-wear. | Slovgaard et al. 2023 [47] |

The definitions, values and the supporting references for the parameters used in the pre-processing operations.
