## Supplementary Figure 3 for "Validation and optimisation of wearable accelerometer data pre-processing for digital measure implementation and development"

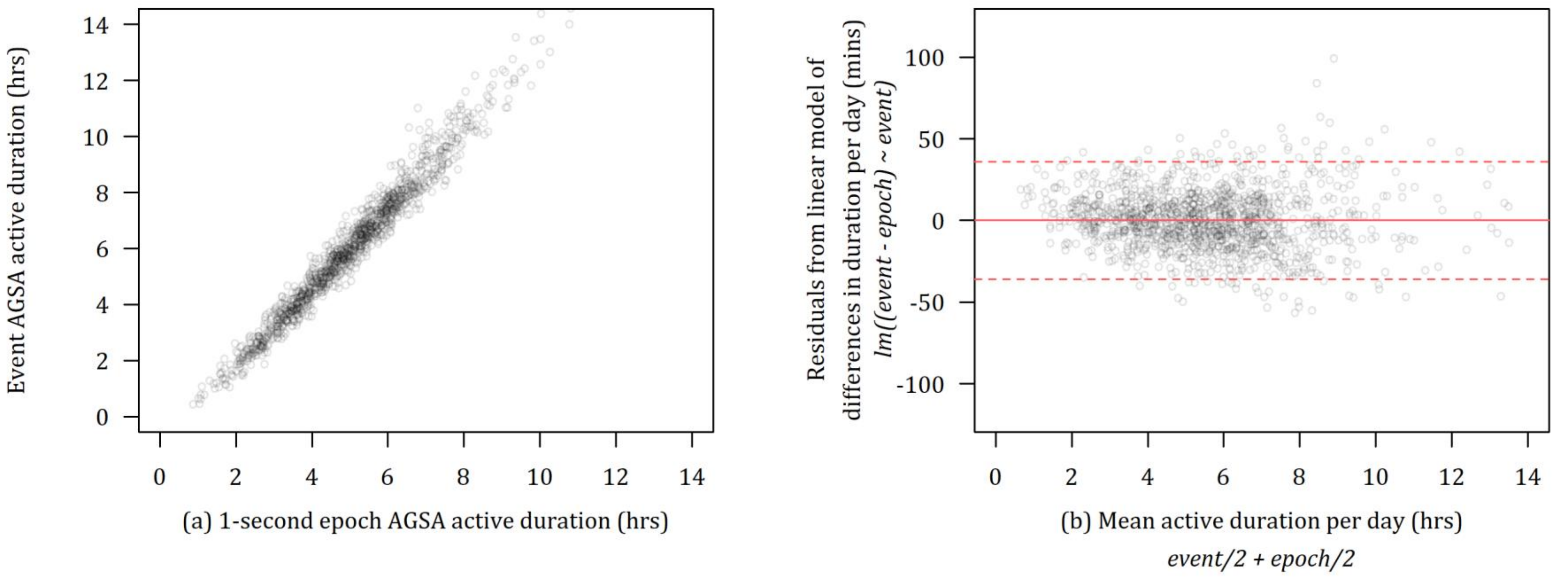


**S3 Fig. Comparing daily active duration using event and epoch aggregates of AGSA.** (a) Correlation plot of active hours per day as assessed by AGSA 1s epoch and variable duration events, (b) Bland-Altman regression plot modelling the differences of AGSA 1s epoch and variable duration event active hours by event-assessed duration per day with mean bias and 95% SEE confidence intervals.
